# Profiling of cytokines, chemokines and growth factors in saliva and gingival crevicular fluid

**DOI:** 10.1101/2021.03.11.434959

**Authors:** Yong Liu, Renping Zhao, Bashar Reda, Wenjuan Yang, Matthias Hannig, Bin Qu

## Abstract

In saliva and gingival crevicular fluid (GCF) soluble factors such as cytokines, chemokines and growth factors have shown a great potential serving as biomarkers for early detection and/or diagnosis of oral and systemic diseases. However, GCF and saliva, which one is a better source is still under debate. This study aimed to gain an overview of cytokines, chemokines and growth factors in saliva and GCF to pave the way for selecting suitable oral fluids for oral and systemic diseases. Multiplex cytokine assay was conducted to determine concentrations of cytokines, chemokines and growth factors in saliva and GCF samples from healthy subjects. The protocol for sample collection was carefully optimized. Stabilization, repeatability, and donor variation of the profiles were analyzed. We found that for different donors, cytokine and chemokine profiles showed unique patterns in saliva but similar patterns in GCF. In terms of growth factors, the profiles were individualized in saliva and GCF. All profiles stayed stable for the same healthy individual. In saliva, profiles of cytokines, chemokines and growth factors are individualized for different donors. In GCF, profiles of cytokines and chemokines are similar. Other factors, such as growth factors and T helper-related cytokines, are highly variable in donors. Profiles of soluble factors are not correlated in saliva and GCF. The comprehensive cytokine profiles in saliva and GCF reported in this work would serve as a good base for choosing promising cytokines for developing biomarkers in oral fluids.

## 1. Introduction

Gingival crevicular fluid (GCF) and saliva are the main fluids found in the oral cavity. Gingival crevicular fluid (GCF) is an exudate of the plasma/serum originating from the blood vessels located in the gingival connective tissue, exuding into the dentogingival space. Saliva is watery liquid mixture secreted by salivary glands into the oral cavity. GCF and saliva play a critical role in maintaining physical functions, protecting oral tissues from pathogens and oral diseases ^1^.

Cytokines are small proteins serving as extracellular messengers, which are involved in regulating cell functions, especially for immune cells ^2^. Emerging evidence shows that certain cytokines in saliva or GCF serve as promising biomarkers to determine oral or systemic diseases. In saliva, early gingival inflammation is associated with a reduction in the level of salivary IL-8, an inflammatory cytokine ^3^. Studies suggest that specific salivary cytokines could be promising biomarkers for an early-stage diagnosis of heterotopic ossification ^4^ and of oral cancer ^5^. In GCF, Studies show that the profiles of cytokines as well as growth factors differ between healthy individuals and patients with periodontal diseases ^6–9^, gingival inflammation, and rheumatoid arthritis ^9^. Among these cytokines/mediators, IL-1α, IL-1β and IL-17a are suggested as promising biomarkers to distinguish patients with chronic periodontitis from healthy individuals ^10^. Despite the promising potential of cytokines in saliva and GCF, cytokine profiles in saliva and GCF have been mostly investigated separately ^7,9,11–14^. A thorough overview and systematic comparison of profiles of these immune-related factors (cytokines, chemokines and growth factors) between saliva and GCF, especially in healthy individuals, is still missing.

In this study, we used the multiplex cytokine assay to analyze concentrations of cytokines and growth factors in saliva and GCF samples from healthy donors. Comparing profiles among different donors, cytokine profiles are comparable in GCF, but individualized in saliva. Concerning growth factors, their profiles exhibited unique patterns in GCF and saliva for each subject.

## 2. Materials and methods

### 2.1 Collection of saliva and GCF samples

Healthy volunteers (14 females and 11 males) aged 21-57 y (median: 30 y) were recruited. For saliva, each individual expectorated around 2 ml of unstimulated saliva into sample tubes. Before GCF collection, the tooth surface was carefully cleaned and soft plaques were removed. Paper strips made from osmometer sample discs (ELITechGroup, USA) were carefully inserted into the gingival culcus of the teeth for 30 seconds until the stripes were fully saturated. The strips contaminated with blood were not included for further analysis. All samples were kept at −80°C until use.

### 2.2 Multiplex cytokine assay

Saliva samples were centrifuged and the supernatant was collected. For GCF samples, 130 μl of assay buffer was added into each tube which contained four paper strips. The supernatant was harvested for the multiplex cytokine assay (LEGENDplex, Biolegend), which was conducted according to manufacturer’s instructions without modifications. Each sample was analyzed in duplicates.

## 3. Results and discussion

### 3.1 Cytokines are stable in GCF and saliva

To investigate cytokine profiles in saliva and GCF, we verified that rinsing the oral cavity with water did not significantly alter the pattern of the cytokine profiles in GCF (Supplementary Fig. 1A, B) and saliva (Supplementary Fig. 1C). In addition, leaving saliva samples at room temperature for 24 hours did not significantly affect cytokine profiles compared to the directly frozen samples (Supplementary Fig. 1D). The results thus indicate that cytokine profiles are considerably stable. Minor variations in sample collection conditions would not lead to drastic change in cytokine profiles. It also demonstrates that this cytokine assay is a robust method, which can reliably and reproducibly determine cytokine concentration.

### 3.2 Cytokine profiles are universal in GCF but individualized in saliva

To gain a thorough overview of cytokine profiles in saliva and GCF, we first recruited four healthy volunteers, from whom samples were collected 3 days in a row. We started with inflammatory cytokines, as pre-set in the Human Inflammation Panel. We found that in GCF samples, the cytokine profiles are very similar among different donors, featuring with a prominent peak for IL-8 (Fig. 1A). Unexpectedly, salivary cytokine profiles showed individualized features (Fig. 1B). In addition, we noticed that for the same donor, the pattern of the cytokine profiles stayed comparable throughout three days in both saliva and GCF samples (Fig. 1A, B). Unexpectedly, for the same donor, GCF and saliva exhibited very distinctive cytokine profiles (compare Fig. 1A and 1B). For different donors, the patterns of cytokine profiles in GCF were similar and in a comparable range, whereas in saliva the difference among donors is very prominent. We examined further healthy individuals (Fig. 1C, D, Supplementary Fig. 2), cytokine profiles of saliva and GCF samples of which supported this conclusion. These are no difference identified in female donors (Donor 1-14) versus male donors (Donor 15-25). These findings indicate that GCF and saliva possess distinctive compositions of inflammatory cytokines, which stay relatively stable with time for the same healthy individual.

**Figure 1.**
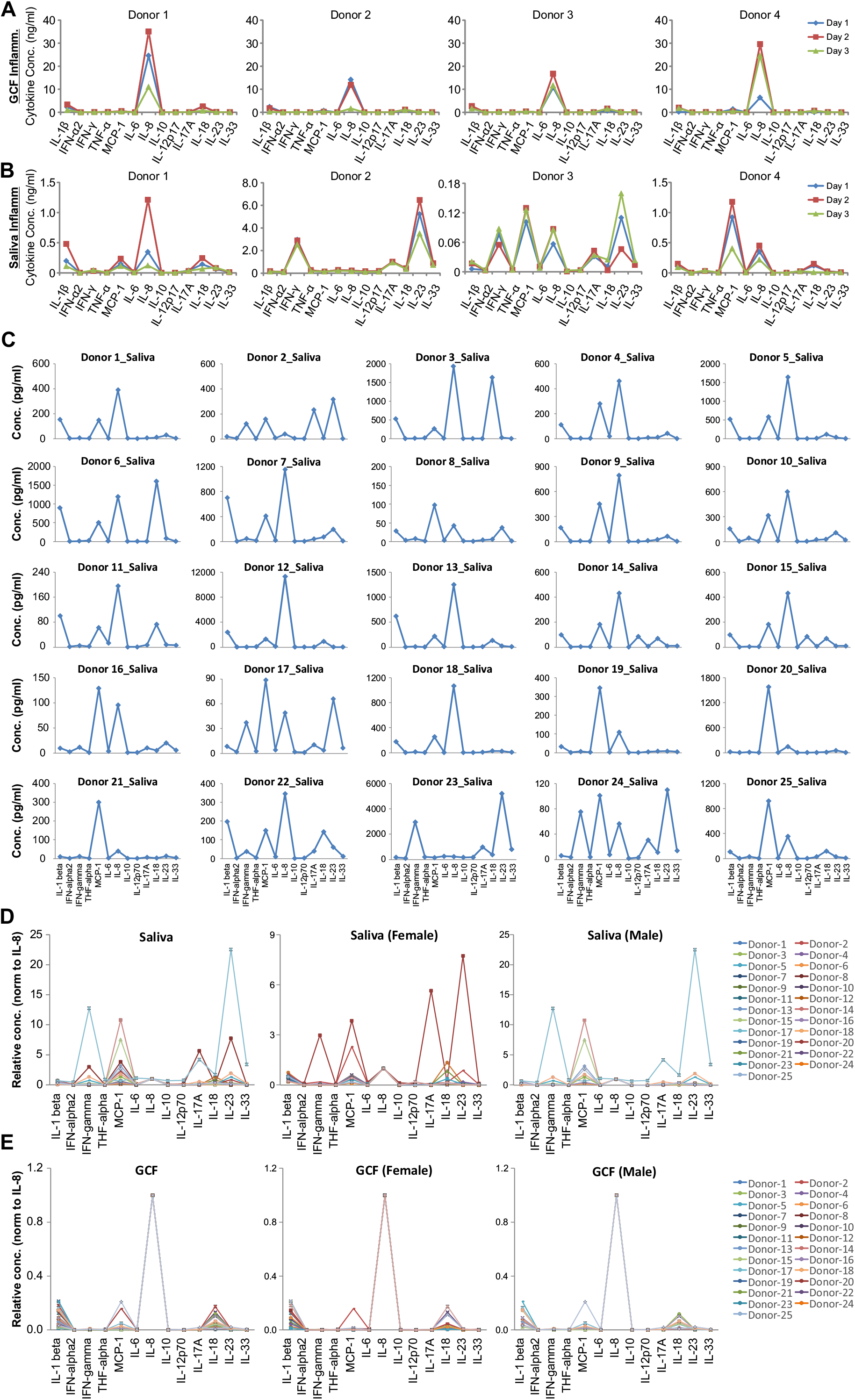
Profiles of inflammatory cytokines in GCF and saliva. Concentration of cytokines was determined using a bead-based multiplex cytokine assay with the pre-set Human Inflammation Panel. (**A, B**) Samples were collected either from four male donors on three consecutive days. (**C-E**) 14 female donors (Donor 1-14) and 11 male donors (Donor 12-25) were recruited.

We further analyzed the profiles of other cytokines, such as pro-inflammatory chemokines (Fig. 2A, B) and the cytokines secreted mainly by macrophages and stromal cells (Fig. 2D, E). We found that, in good agreement with inflammatory cytokines, for different donors the profiles of these cytokines in GCF are similar (Fig. 2A, C). In comparison, the salivary profiles of these two panals were very different (Fig. 2B, D). Overall, these patterns of cytokine profiles in saliva and GCF were not much altered with time for each individual (Fig. 2A-D). Together, our findings show that cytokine profiles are, in general, individualized in saliva whereas common features are share in cytokine profiles in GCF.

**Figure 2.**
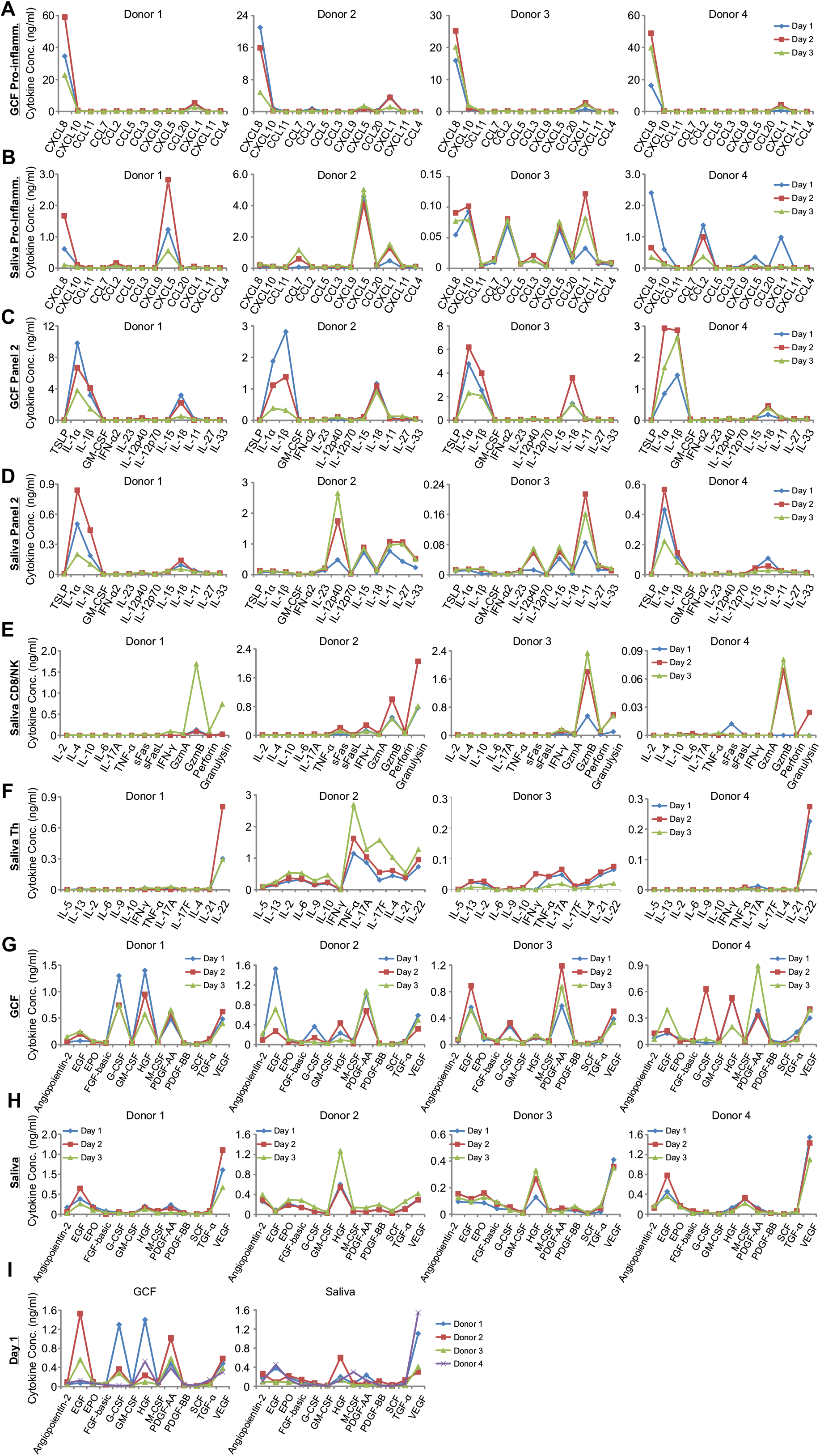
Profiles of other soluble factors in saliva and GCF. Concentration of the factors was determined using bead-based multiplex cytokine assay with the pre-set Human Pre-inflammation cytokine Panel (**A, B**), Panel 2 (**C, D**), CD8/NK Panel (**E**), Th Panel (**F**), or Growth Factor Panel (**G-I**).

Immune killer cells, namely CD8^+^ T cells and natural killer (NK) cells play a pivotal role to eliminate pathogen-infected or tumorigenic cells. T helper cells (Th) act as a coordinator to orchestrate immune responses. Therefore, we also analyzed the cytokines and cytotoxic proteins, which are involved in the effector function of immune killer cells (CD8/NK) or T helper cellls (Th). In saliva, a significant amount of the cytotoxic proteins granzyme B (GzmB) and granulysin were detected, which could vary with time (Fig. 2E). The cytokines involved in Th functions, showed highly variable expression among different donors (Fig. 2F, Supplementary Fig. 3A). In GCF samples, the concentrations of Th-cytokines are much lower than inflammation and pro-inflammation cytokines; the profiles are not always similar among different donors (Supplementary Fig. 3B).

### 3.3 Profiles of growth factors vary significantly with time and are individualized for GCF and saliva

At last, we analyzed the profiles of growth factors in GCF and saliva. We found that in GCF, the profiles of growth factors are not alike among donors (Fig. 2G). Within three days, the shape of the growth factor profiles stayed relatively similar for Donor 1 and Donor 3, but remarkably changed for Donor 2 and Donor 4 (Fig. 2H). In saliva, the profiles of growth factors were distinctive for all four donors and exhibited not much change within the time period examined (Fig. 2H). The concentration of growth factors was found, in most cases, higher in GCF compared to saliva (Fig. 2I). These results suggest that the profiles of growth factors in GCF are individualized and could be changed with time.

In summary, our study shows that profiles of cytokines, chemokines and growth factors differ between saliva and GCF. For different donors, the patterns of cytokine and chemokine profiles are personalized in saliva but similar in GCF. In terms of growth factors, the profiles diverge in saliva and GCF. In the same healthy individual, the profiles of soluble factors are relatively stable. In general, GCF has less extend of variation among different donors, making it a good candidate for searching biomarkers.

## 4. Acknowledgements

We thank the donors for providing GCF and saliva samples. We thank Markus Hoth (Saarland University) for the valuable inputs and for the use of flow cytometer. The flow cytometer was funded by DFG (GZ: INST 256/423-1 FUGG). The work was supported by German Research Foundation (Sonderforschungs-bereich (SFB) 1027 project A2 (to B.Q.), project B3 (to M.H.), miniproposal of SFB1027 (to Y.L.)).

## 6. Ethical considerations

All procedures performed in studies involving human participants were in accordance with the ethical standards of the institutional and/or national research committee and with the 1964 Helsinki declaration and its later amendments or comparable ethical standards. The study protocol has been approved by the Medical Ethics Committee of the Medical Association of Saarland, Germany (# 238/03, 2016, 2020). Informed consent was obtained from all individual participants included in the study.

## Supplementary Information

**Sup.Figure 1.**
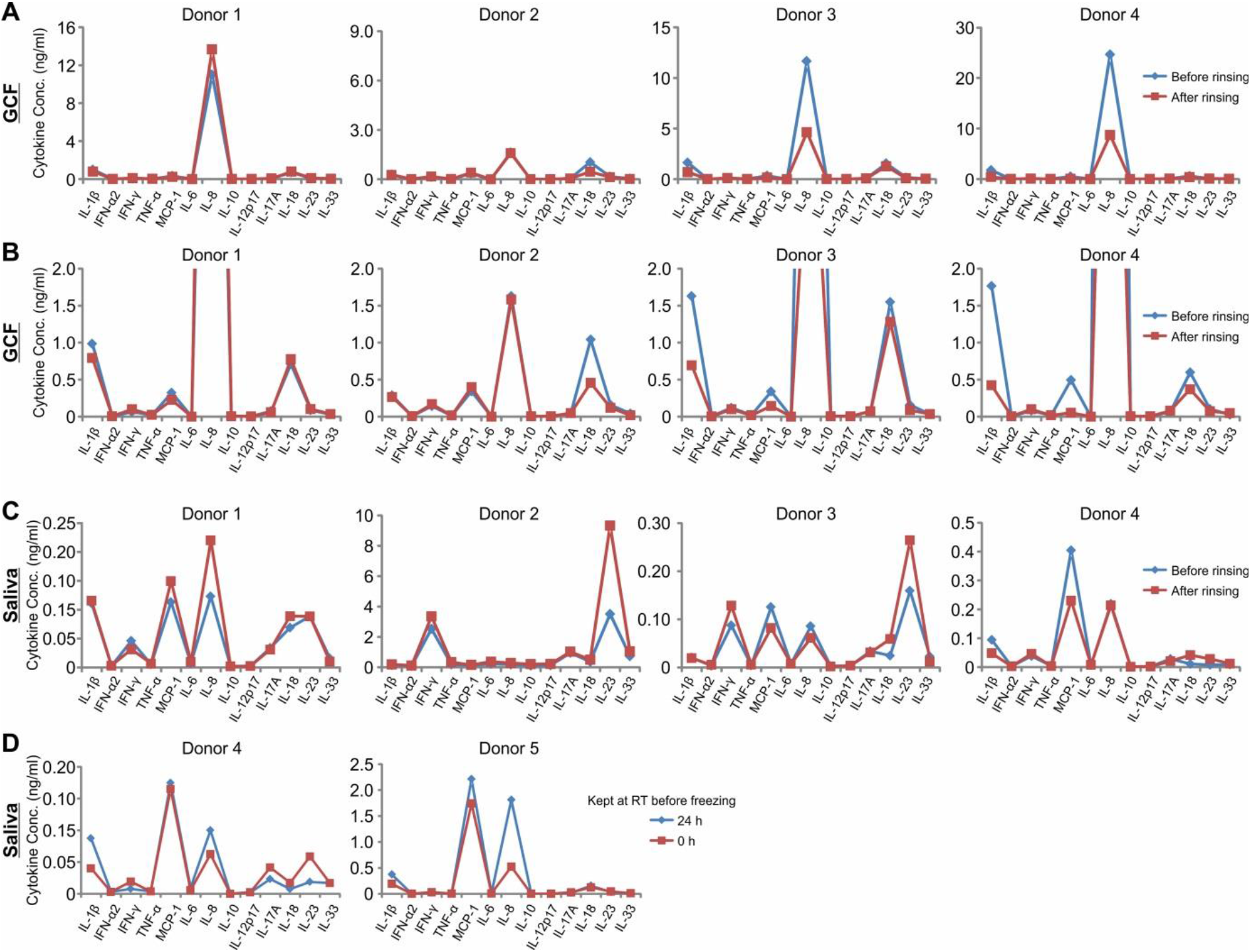
Cytokines are stable in GCF and saliva samples. The volunteers rinsed their oral cavity with 100 ml ddH_2_O for 30 seconds. Unstimulated saliva and GCF were sampled after 2 minutes. The profiles of inflammatory cytokines were determined using the multiplex cytokine assay. **A-C.** Cytokine profiles can be quickly recovered from water rinsing in GCF (A, up to 30 ng/ml; B, 0 - 2 ng/ml) and in saliva (C). **D.** Cytokine profiles are stable at room temperature for 24 hours.

**Sup.Figure 2.**
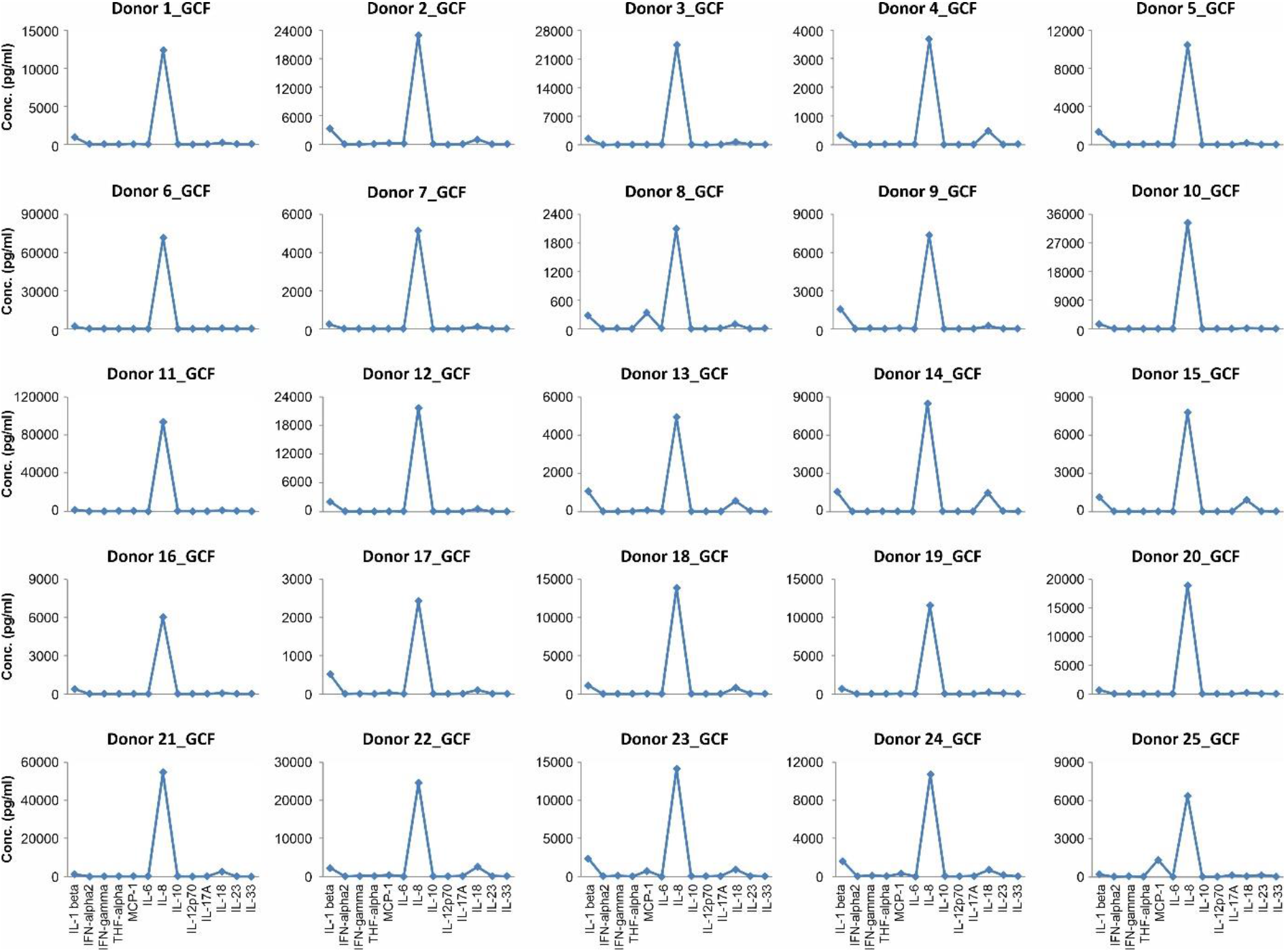
Profiles of inflammation cytokines in GCF. GCF samples were collected as described in the Methods. Concentration of cytokines was determined with the pre-set Human Inflammation Panel using a bead-based multiplex cytokine assay.

**Sup.Figure 3.**
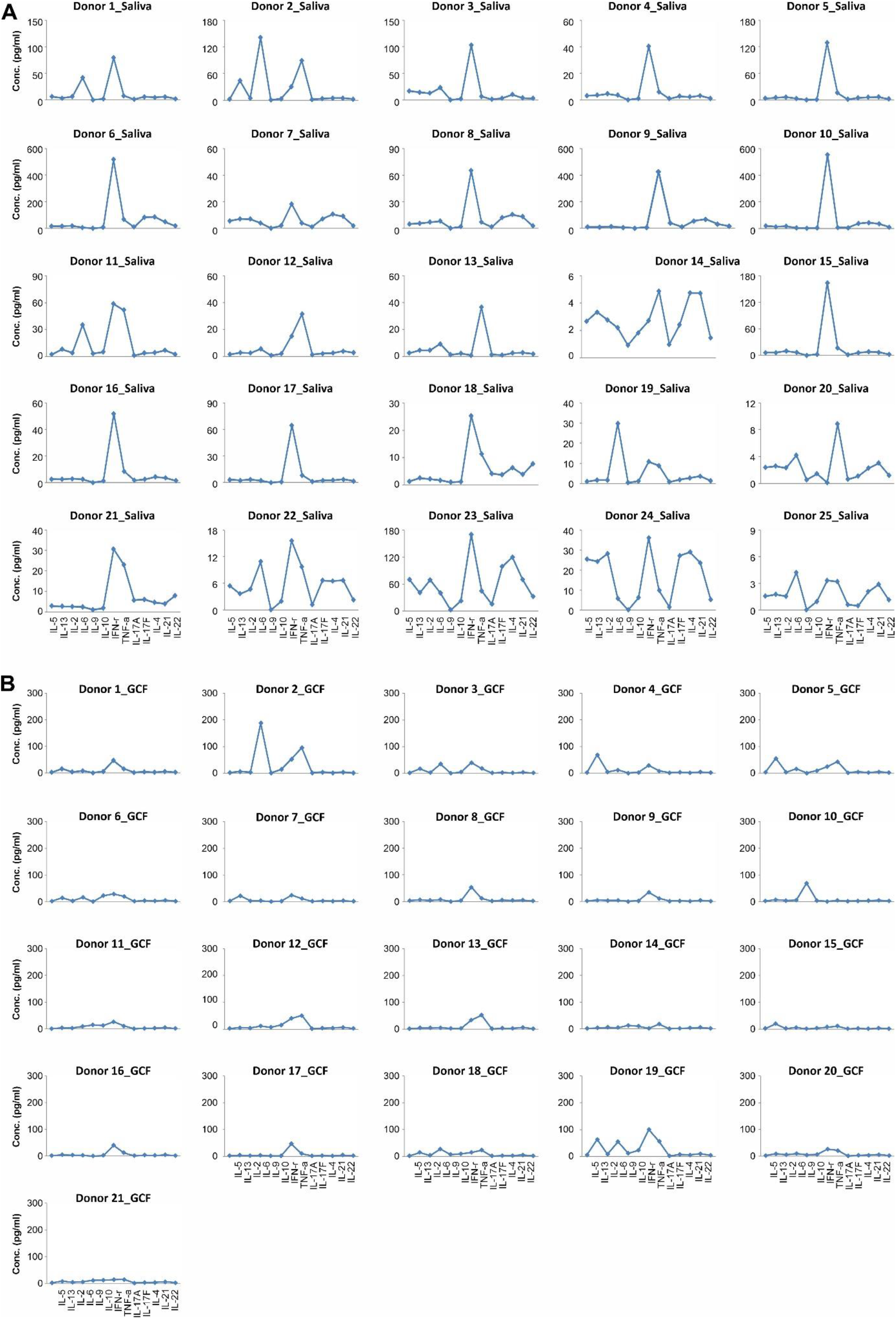
Profiles of Th-related cytokines in GCF and saliva. Saliva (A) and GCF (B) samples were collected as described in the Methods. Concentration of cytokines was determined with the pre-set human Th Panel using a bead-based multiplex cytokine assay.

